# Identification of Novel and Differentially Expressed MicroRNAs in Goat Enzootic Nasal Adenocarcinoma

**DOI:** 10.1101/057885

**Authors:** Bin Wang, Ni Ye, San-jie Cao, Xin-tian Wen, Yongxs Huang, Qi-gui Yan

## Abstract

Enzootic nasal adenocarcinoma (ENA), an epithelial tumor induced in goats and sheep by enzootic nasal tumor virus (ENTV), is a chronic, progressive,contact transmitted disease. This study aimed to identify novel and differentially expressed miRNAs in the tumor and para-carcinoma nasal tissuesofNanjiang yellow goats with ENA. Small RNA Illumina high-throughput sequencing was used to construct a goat nasal miRNA library.

406 known miRNAs and 29 novel miRNAs were identified. A total of 116 miRNAs were significantly differentially expressed in para-carcinoma nasal tissues and ENA; Target gene prediction and functional analysis revealed that 6176 non-redundancy target genes, 1792 significant GO and 97 significant KEGG pathway for 121 miRNAs were predicted.

---

MicroRNAs (miRNAs) are endogenous, 21-24 nucleotide-long, non-coding RNAs that regulate gene expression in eukaryotes; however, some viruses alsoexpress miRNAs in host cells (Bartel 2004; Pfeffer et al. 2004; Filipowicz et al. 2005). MiRNAs are complementary to specific sequence motifs in the 3’ untranslated regions (UTRs) of their target mRNAs and negatively regulate gene expression at the post-transcriptional level by inhibiting translation or promoting mRNA degradation, based on the degree of complementary base pairing between the miRNA and mRNA. MiRNAs regulate approximately 30% of genes in higher eukaryotic cells, including genes involved in development, metabolism, apoptosis, proliferation and viral defense(Kincaid and Sullivan 2012; Dong et al. 2013; Zhang et al. 2013; Wang and Kaufman 2014; Zhang et al. 2014; Bandiera et al. 2015; Lin et al. 2015). Theearliest evidence for an association between miRNAs and cancer came from the study of chronic lymphocytic leukemia (CLL) (Calin et al. 2002). To date, more than 50% of miRNAs have been shown to be encoded in chromosome fragile sites that are often absent, amplified or rearranged in malignant tumor cells leading to dysregulated expression of miRNAs, and numerousmiRNAs have been shown to play important roles in tumorigenesis (Calin et al. 2004). MiRNAs can act in a similar manner to oncogenes or tumor suppressor genes and have emerged as a novel type of regulatory factor in the epigenetic modification of gene expression. According to predictions in vertebrates, a single miRNA can regulate more than 400 target genes, forming complicated regulatory networks(Lewis et al. 2003; Landgraf et al. 2007; Ruby et al. 2007; Bartel 2009). Therefore, miRNAs have become a focus of cancer research in order to identify novel molecular methods for the diagnosis, prognostication and treatment of human cancer. Now researchers can directly obtain miRNA sequences and discover novel miRNAs through utilize Illumina high-throughput sequencing technology(Ye et al. 2012).

Enzootic nasal adenocarcinoma (ENA) is an epithelial tumor caused by enzootic nasal tumor virus (ENTV), and is a chronic, progressive, contact transmitted disease(Heras et al. 2003). With the exception of Australia and New Zealand, this disease has spread throughout goats or sheep almost worldwide (Kawasako et al. 2005). ENA originates from the ethmoid area of the nasal cavity either unilaterally or bilaterally, and the tumors are soft, whitish or pinkish-red in color and can partially or completely obscure the nasal cavity(Walsh et al. 2010). Metastases to the regional lymph nodes,brain or other organs does not occur(Yi et al. 2010). So far, there are no effective methods for early diagnosis of ENA and the goats or sheep can only be culled after symptoms appear. More seriously, as it is difficult to distinguish between animals with a latent infection and healthy animals, the virus spreads within herds, and can infect a large number of goats or sheep and threaten the entire population.

There are currently 2581 human miRNAs in the miRbase (v21) database; however, there is no public miRNA library of *Capra hircus* nasal tissues and there have been no reports of miRNAs in ENA. To further complicate matters, attempts to establish a system of cultivating ENTV *in vitro* have failed, which presents a significant obstacle to investigating the immunological characteristics of ENTV and the mechanisms by which it promotes tumorigenesis (De las Heras et al. 1995). Therefore, taking advantage of knowledge ofthe roles of miRNAs in human cancer to research the miRNAs involved in ENA may not only avoid the problem of cultivating ENTV *in vitro*, but also shifts the focus to the cells targeted by ENTV - goat or sheep nasal epithelial cells - and may providean alternative method for investigating the tumorigenic effects of ENTV.

Using Illumina high-throughput sequencing technology to detect miRNAs expressed in the tumor and para-carcinoma nasal tissues of Nanjiang yellow goats with ENA, we constructed the first goat nasal tissue miRNA library. Furthermore, the target genes of the differentially expressed miRNAs in ENA were predicted and their corresponding biological functions were analyzed. This research may help to identify novel biomarkers for ENA, lays a foundation for investigating the mechanism by which ENTV promotes tumorigenesis, and provides further information on the role of miRNAs in cancer. Furthermore, as the sequences and roles of miRNAs are well conserved, the findings of this study may also be relevant to human cancers such as nasopharyngeal carcinoma.

## RESULTS

### Capra hircus nasal tissue miRNA library

High-throughput sequencing generated hundreds of millions of reads for each tissue. The raw data (tag sequences and counts) have been submitted to Gene Expression Omnibus (GEO) under series GSE65305. To estimate sequencing quality, the quality scores were analyzed across all bases (Fig. 7). The lowest quality score was ≥ 30; therefore, the error rate was lowerthan 0.1%. Reads including adaptor sequences, low quality sequencesand sequences of unqualified length were removed, and the remaining clean reads were aligned with the *Capra hircus* genome in NCBI using Bowtie software to analyze the genomic distribution and expression ofsmall RNAs. The vast majority of clean reads (at least 84.75%) and unique reads (at least 57.56%) mapped to the *Capra hircus* genome (Table 1).

**Table 1.**
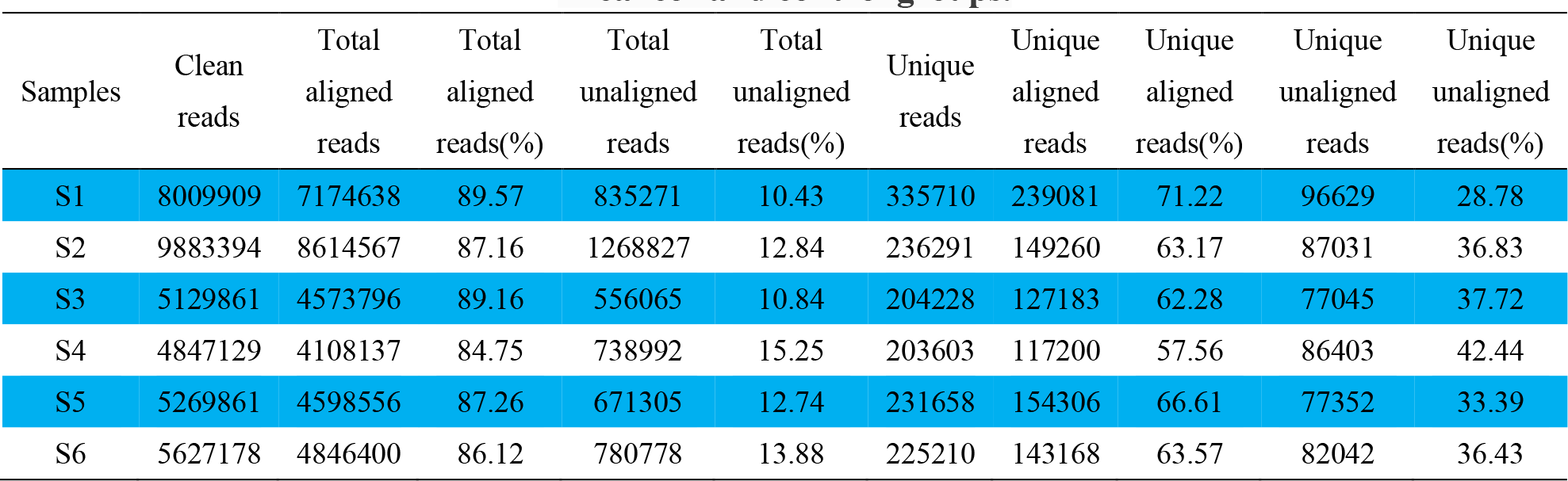
The results of clean reads and unique reads maped to the *CapraHircus* genome in cancer and control groups.

Unique reads were blasted against the Rfam, RepBase, EST and miRBase databases, in the order of known miRNAs > rRNAs > tRNAs > snRNAs > snoRNAs > repeat sequences, which enabled each small RNA obtain a unique annotation. To exclude other RNAs, such as tRNAs, rRNAs, snRNAs and snoRNAs, the remaining sequences were mapped to Denovo predictiondata sets and the *Capra hirus* genome to identify novel miRNAs. Using miRDeep prediction (Friedländer et al. 2008) and RNAfold (Hofacker et al. 1994) software to analyze secondary structure. A total of 435 sequences (29 novel miRNAs) were included in the miRNA library. Table S1 displays the sequencing generated codes and corresponding *Capra hirus* miRNAs or novel miRNA_id. As research into miRNAs in human cancer is widespread, we blasted all goat miRNAs against the human miRNAs in miRBase v21 to further understand their function. A total of 615 of the goat miRNAs had analogues in the human miRNA datasets (Table S2).

### Identification of differentially expressed miRNAs in ENA

A total of 435 miRNAs were identified in the ENA and para-carcinoma tissues. The table S3 lists the expression of all miRNAs. The expression of 116 miRNAs was significantly different in para-carcinoma tissues and ENA, of which 54 were downregulated and 60 were upregulated in ENA. In addition, 2 miRNAs were only expressed in the para-carcinoma tissues (Table S4). The majority of the fold change-log^2^ values ranged from 1 to 5.42; chi-miR-133a-3p had the highest fold-change-log^2^ of at least 5.4-fold, and 65 miRNAs had fold-change-log^2^ values of at least two-fold. Figure 8 indicates the differences in expression of all 435 miRNAs between ENA and the para-carcinoma tissues.

### Functional analysis of differentially expressed target genes regulated by differentially expressed miRNAs

The Miranda algorithm indicated thousands of potential target genes forthe 435 miRNAs. According to the total scores and predicted energies, the 6176 non-redundancy target genes of these 121 miRNA(116 significant expression miRNAs and 5 star miRNAs) were selected, reflecting 15222 corresponding relationships between the differentially expressed miRNAs and their target genes. Table S5 shows the total stores, total energy, and protein-id and genomic location of predicted target genes.

The expression of these candidate target genes was assessed in the high throughput sequencing data obtained from the same ENA and para-cancerous tissue samples (data not shown; this data will be described in another article). A total of 175 mRNAs that were significantly differently expressed in ENA were selected for this analysis. Table S6 lists the differentially expressed miRNAs and their corresponding differentially expressed target genes. Table S7 summarizes the degree of regulation between the differentially expressed miRNAs and their differentially expressed target mRNAs.

### MiRNA-gene ontology network analysis of miRNA target genes

Functional analysis was conducted on the mRNAs predicted as targets of the 435 differentially expressed miRNAs. A total of 9777 GO enrichments were identified, of which 1792 GO categories were significant (*P* ≤ 0.05).The mRNA sequencing identified a total of 90 target genes corresponding to miRNAs with significantly decreased expression and 84 target genes corresponding to miRNA with significantly increased expression in tumor group. 472 significant GO enrichments exist in significant expression miRNA-mRNA network(Table S8). The target genes of the differentially expressed miRNAs were mainly involved in cell may be implicated in the tumorigenic effect of ENTV. Figure 9 presents the ten most-enriched GO categories for the differentially expressed target genes of the differentially expressed miRNAs in ENA.

### Analysis of signaling pathways regulated by the miRNA target genes

Signal transduction analysis was conducted on the mRNAs predicted as targets of the 435 differentially expressed miRNAs, and 267 KEGG enrichments were identified of which 97 were significant (*P* ≤ 0.05). The target genes of the differentially expressed miRNAs participate in pathways related to the signal transduction, specific types of cancer and immune system. Among significantly differently expressed miRNA, miRNA with increased expression in tumor group were involved in 83 significant signal transduction pathways (Table S9), miRNA with reduced expression in tumor group were involved in 89 significant signal transduction pathways (Table S10). Figure 10 illustrates the ten most-enriched KEGG pathways for the differentially expressed target genes of the differentially expressed miRNAs in ENA.

### Quantitative RT-PCR validation of differentially expressed miRNAs

We selected 9 of the key miRNAs that were significantly differently expressed in ENA including five miRNAs that featured in both the GO and KEGG pathway analyses and two novel miRNAs(NW_005102245.1_1433,NC_022308.1_285). The main functions of target genes regulated by these miRNAs are involved in cancer pathogenesis, virus infection, cell apoptosis and proliferation. As shown in Figure 11, qRT-PCR confirmed the expression of the nine miRNAs between ENA and the para-cancerous tissues with an increased sample size. The expression trend of eight miRNAs is in accord with Illumina High-Throughput Sequencing, one miRNA(chi-miR-218) both in sequencing and qPCR verification have shown a down-expression in tumor group, but the down-expression multiple is different.

## Discussion

ENTV, a betaretrovirus that infects sheep (ENTV-1) and goats (ENTV-2), is associated with neoplastic transformation of ethmoid turbinate epithelial cells and leads to ENA. The clinical symptoms of goats are a loss of appetite, extreme weight loss, dyspnea, rhinorrhea, and unilateral or bilateral nasal puffiness. The incidence of ENTV infection ranges from 5% to 15%, and once the clinical symptoms of ENA appear, almost all cases are fatal (Rings and Rojko 1985; De las Heras et al. 1991; Vitellozzi et al. 1993). High-throughput sequencing technology is gradually being used in animal and has provided some knowledge of goat miRNAs. Ji et al. (Ji et al. 2012) discovered 290 known miRNAs and 38 novel miRNAs in dairy goat mammary gland tissue and reported that miRNA-mediated regulation of geneexpression occurs during early lactation. Hao et al. (Yu and Jun 2014)found that the expression of 64miRNAs was reduced in the skin of a 70-day fetus relative to a lamb born at 2 weeks, with the expression of ten miRNAs decreasing more than 5-fold, which implies that miRNAs play an important role in maintaining normal skin function.

Cancer is a leading cause of morbidity and death in humans. Significantresearch has been conducted on miRNAs in human cancer, and miRNAs have been demonstrated to be directly involved in human nasopharyngeal carcinoma (NPC). For example, miR-29c, the miR-34 family, miR-143, miR-145 and miR-9 are downregulated in NPC, leading to increased expression of their target genes which influence the function and synthesis of extracellular matrix proteins, which in turn affects tumor invasion and metastasis, and activates the TGF-Wnt, IP3 and VEGF signaling pathways (Sengupta et al. 2008; Chen et al. 2009). In contrast, miR-200, the miR-17-92 cluster and miR-155 are upregulated in NPC, and miR-200 inhibits the migration and invasion of NPC cells by inhibiting the expression of *ZEB2* (zinc finger E-box binding homeobox 2) and *CTNNB1*(catenin-β-like 1) (Xia et al. 2010). Byblasting the 435 miRNAs identified using high-throughput sequencing in this study against the human miRNA datasets in miRBase, we found that hsa-miR-9, hsa-miR-34 and hsa-miR-143 are significantly downregulated and hsa-miR-200 is significantly upregulated in ENA. GO and KEGG pathway analysis revealed these miRNAs are involvedin intracellular signal transduction, the MAPK cascade and cell morphogenesis, among other processes.

Our study found according to the percentage, the top five signaling pathways are MAPK signaling pathway Pathways in cancer PI3K-Akt signaling pathway Ras signaling pathway and Viral carcinogenesis. Kano et al. (Kano et al. 2010) and Chiyomaru et al. (Chiyomaru et al. 2010) found that miR-133awas significantly inhibited human esophageal squamous cell cancer and the invasion of bladder cancer cell. Iorio (Iorio et al. 2005) found that expression of miR-133a significantly reduced during the progression of breast cancer. Our results also reveal the expression of miR-133a-3p was at least5-fold lower in ENA compared to para-carcinoma nasal tissues. These results suggest that miR-133a-3p may regulate the expression of oncogenes and inhibit tumorigenesis. KEGG analysis displayed that the target genes of miR-133a-3p are involved in tumor biology at multiple nodes, such as regulation of cell differentiation, apoptosis, signal transduction and cell adhesion, invasion and migration. In esophageal squamous cell carcinoma and bladder cancer, miR-133a targets fascin actin-bundling protein 1(*FSCN1*) to regulate cancer cell invasion, migration and proliferation (Iorio et al. 2005; Kano et al. 2010). However, in this study we identified that serine/threonine-protein kinase B-raf(*BRAF*)as chi-miR-133a-3p, chi-miR-145-5p,chi-miR-146a/200a and two novel miRNA(NC-0223 08.1-260-NC-022294.1-874) target gene which acts upstream regulatory factor in RAS-RAF-MEK-ERK. Sustained activation of BRAF will lead to cell deterioration and excessive proliferation(Ji et al. 2007). In addition, the miR-133a-3p target genes:*MDS1* and *EVI1* complex(*MECOM*)may also play a significant role in pathways related to cancer. In chronic myeloid leukemia (*CML*), expression of the oncogene *MECOM* correlates with progression. The tyrosine kinase catalytic activity of the oncoprotein BCL-ABL1 regulates *MECOM* expression, and conversely MECOM partially mediates BCR-ABL1 activity (Roy et al. 2012); BCR-ABL1 activates the PI3K, MAPK and JAK-STAT signal transduction pathways (Pendergast et al. 1993; Carlesso et al. 1996; Skorski et al. 1997) to promote abnormal proliferation, differentiation, transformation and survival in myeloid cells (Smith et al. 2003). However, forkhead box O *(FoxO)* as the intersection of PI3K and RAS signaling pathway can inhibit cell proliferation and induce cell cycle stop. The activation of the PI3K signaling pathway inhibits the activity of the FoxO transcription factor(Schmidt et al. 2002; Martínez-Gac et al. 2004),which increase the chances of tumor formation. Further study is required to determine if *MECOM* and BCR-ABL1 play a role in the pathogenesis of ENA.

miR-148a is an oncogene that is upregulated in hepatocellular carcinoma cells(*HCC*)and enhances cell proliferation, migration, invasion and stimulates the epithelial to mesenchymal transition *(EMT)* by targeting tumor suppressor gene: phosphatase and tensin homolog(*PTEN*)(Yuan et al. 2012). However, *PTEN*was not identified as a target of miR-148a in this study. Its predicted targets were the transforming growth factor β receptor associated protein 1 (*TGFβRAP1*)which can specifically combine with the receptor of transforming growth factor β(TGFβ,and then help to realize the biological function of TGFβ(Chen et al. 1995).TGFβ can inhibit cells growth in malignant tumor such as head and neck squamous cancer, colon cancer, breast cancer(Arteaga et al. 1990; Wu et al. 1992; Briskin et al. 1995).The present studies have pointed out that the expression of TGFβ in nasopharyngeal phosphorus tumor generally weakened or even disappear, but the adjacent epithelium have stronger expression(Bin et al. 1999). The expression of miR-148a-3p was at least 2.5-fold higher in ENA compared to para-carcinoma nasal tissues. Influenced by miR-148-3p expression, *TGFβRAP1* will drop, whichwillaffect the signal pathway of TGFβ and make the cancer cell reduction or loss of ability to react to TGFβ, finally, the tumor cells escape from negative growth regulation of TGFβ.Although this is our speculation, but we believe there is a link between them.

## Conclusions

This study provides a solid basis for further research and highlights anumber of miRNAs and genes that may be involved in the pathogenesis of ENA. This study of miRNAs in ENA may also provide useful information for basic research into human cancer. In future studies, we aim to confirm the function of the candidate miRNAs in nasal cells. In addition, we hope that these studies may provide some clues to help establish a method for cultivating ENTV *in vitro*.

## Methods

### Ethics Statement

This study was carried out in strict accordance with the Guidelines forExperimental Animals of the Ministry of Science and Technology (revised in 2004; Beijing, China) and was approved by the Institutional Animal Care and Use Ethics Committee of Sichuan Agricultural University, NO. SYXK (Chuan) 2014-187.

### Animals and tissue samples

Eight goats (3a-4a)infected by ENTV under natural conditions at a farm in Sichuan were quarantined and transported to Sichuan Agricultural University laboratory animal center, and grew up the center. After slaughter, tumor and para-carcinoma nasal tissues were collected, frozen rapidly in liquid nitrogen and stored at −80°C. After pathological analysis, the samples from three Nanjiang yellow goats whose nasal passages were unilaterally blocked by tumors were selected for high-throughput sequencing. The nasal tumors in these animals were poorly differentiated (i.e., at the same state of differentiation) with no tumor cell infiltration in the matchedpara-carcinoma tissues.

### Preparation of samples for sequencing and qPCR

Samples of cDNA from the tumor tissues (numbers S1, S3, S5) and matchedpara-carcinoma tissues (numbers S2, S4, S6) of the three animals described above were shipped on dry ice to Jing Neng Bio-Technology corporation (Shanghai, China) for high-throughput sequencing. Briefly, total RNA was extracted from the tissues using RNAzol RT RNA Isolation Reagent (GeneCopoela, Rockville, MD, USA) according to the manufacturer’s protocol. The RNA concentrations were determined using a Smart Specplus Spectrophotometer (Bio-Rad, Hercules, CA, USA) and the integrity of the total RNA samples was verified by polyacrylamide gel electrophoresis (PAGE). The All-In-One miRNA qRT-PCR Detection Kit (GeneCopoela) was used to add poly(A) tails to the miRNAs in the total RNA samples and M-MLV reverse transcriptase was used to synthesize cDNA according to the manufacturer’s instructions. Each reaction mixture contained 5 μL of 5x reaction buffer, 1μL RTase Mix, 1 μL of 2.5 U/ μL PolyA Polymerase, 2 μg total RNA and RNase-/DNase-free H2O to 25 μL, and was incubated at 37 °C for 1 h and then at 85°C for 5 min to inactivate the enzyme.

### Analysis of sequence data and creation of miRNA library

Single-read 50 bp sequencing was adopted for high-throughput sequencing. Illumina CASAVA software was used to convert the original data image files into sequence files, and FastQC statistical software was used to evaluate the quality of the data. Primer, adaptor and low quality sequences wereexcluded and 15-40 base sequences meeting the length and quality requirements were selected as clean reads of reliable quality for further analysis(Figure 1-6). Total clean reads from each individual sample were aligned with the Capra hircus genome in NCBI (ftp://ftp.ncbi.nlm.nih.gov/genomes/Capra_hircus) using Bowtie software (Langmead et al. 2009)(http://bowtie-bio.sourceforge.net/index.shtml), and then blasted against the Rfam (http://www.sanger.ac.uk/resources/databases/rfam.html), RepBase (http://www.girinst.org/repbase/), EST (http://www.ncbi.nlm.nih.gov/nucest/) and miRBase (http://www.mirbase.org/) databases. Sequence alignment was set to allow only a single base mismatch, and the results were sorted in the order of known miRNAs > rRNAs > tRNAs > snRNAs > snoRNAs > repeat, respectively, which enabled each small RNAto obtain a unique annotation. The remaining sequences were mapped to Denovo prediction data sets (Bonneau et al. 2006)and the *Capra hirus* genome to exclude known non-miRNA sequences (such as tRNAs, rRNAs, snRNAs and snoRNAs) and identify novel miRNAs. MiRDeep (Friedländer et al. 2008)and RNAfold (Hofacker et al. 1994)were used to predict miRNA precursor sequences, star miRNAs and mature miRNAs, and then the energetic stability, position and read frequencies for each potential miRNA precursor were computed using miRDeep according to the compatibility of energetic stability, positions, frequencies of reads. Ultimately, a *Capra hirus* nasal tissue miRNA library was created by combining the sequencing data from all six samples.

### Identification of differentially expressed miRNAs in ENA

The sequences in each sample were compared with the miRNA library established in this study by assessing the numbers of transcripts per million (TPM). TPM was calculated as (numbers of each miRNA matched to total reads)/(number total reads) × 10^6^. TPM is an indicator of the quantity of miRNA expression per million match paired sequences. The total numbers of matched pair reads were used in the normalized numerical expression algorithm to calculate miRNA expression. DESeq (Anders and Huber 2010)software was used to identify differentially expressed miRNAs between the para-carcinoma tissues (S2, S4, S6) and ENA (S1, S3, S5) on the basis of a fold-change greater than or equal to two and *P*-value ≤ 0.05.

### Prediction and analysis of miRNA target genes

The Miranda algorithm (Enright et al. 2004)was used to predict the target genes of the miRNAs that were differently expressed in ENA. The threshold parameters for predicting miRNA target genes were a total score ≥ 150, ΔG ≤ −30 kcal/mol, and strict 5’ seed pairing. The pathways these candidate target genes are involved in was analyzed by functional annotation utilizing the NCBI, KEGG(http://www.genomejp/kegg/)(Kanehisa et al. 2011) and GO(http://geneontology.org/) (Carbon et al. 2009) databases. Additionally, high-throughput sequencing allowed the mRNA expression of all of the potential target genes to be analyzed in the same samples (data not shown); therefore, GO and KEGG analyses could be conducted on the differentially expressed target genes of the differentially expressed miRNAs. GO annotation and enrichment analysis was performed for three gene ontologies: molecular function, cellular components and biological processes. The following formula was used to calculate the *P*-values:

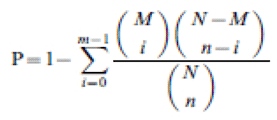

where *N* is the number of genes with GO/KEGG annotations; *n* is the number of target gene candidates in *N*; *M* is the number of genes that annotated to a certain GO term/pathway, and *m* is the number gof target gene candidates in *M*. GO terms and KEGG pathways with a corrected *P*-value ≤ 0.5 were regarded as significantly enriched.

### Validation of the expression of key differentially expressed miRNAs

Key miRNAs that were identified in all of the analyses described above were quantified in ENA and para-carcinoma tissue samples from five goats with ENA whose nasal passages were unilaterally blocked by tumors. Total RNA was isolated and reverse transcribed as described above, then the cDNA products were diluted 5-fold with sterile H_2_O and subjected to quantitativereal-time PCR (qPCR) using the All-In-One miRNA qRT-PCR Detection Kit (GeneCopoela) with U6 snRNA and *GAPDH* as internal references. Each reaction contained 10 μL of 2x All-in-One qPCR Mix, 2 μL All-in-One miRNA qPCR Primer (2 μM; prepared by Life Technologies, Shanghai, China), 2 μL Universal Adaptor qPCR Primer (2 μM), 2 μL first-strand cDNA and 4 μL double distilled water. The cycling conditions were 95 °C for 10 min, 40 cycles of 95°C for 10 s, 60 °C for 20 s and 72 °C for 20 s, followed by melting curve analysis. Relative quantification was performed using the 2^-^ΔΔ^Ct^ method (Livak and Schmittgen 2001), and Mests were used to examine the significance of the differences in expression between the para-carcinoma tissues and ENA.

## Availability of data and material

The raw data (tag sequences and counts) have been submitted to Gene Expression Omnibus (GEO) under series GSE65305.

## Abbreviations

**miRNAs**: MicroRNAs **ENA**: Enzootic nasal adenocarcinoma **ENTV**: Enzootic nasal tumor virus **GO**: Gene Ontology **KEGG**: Kyoto Encyclopedia of Genes and Genomes **RNA**: ribonucleic acid **UTRs**: untranslated regions **CLL**: chronic lymphocytic leukemia **PAGE**: polyacrylamide gel electrophoresis **rRNAs**: ribosomal RNAs **tRNAs**: transfer RNAs **snRNAs**: Small nuclear RNAs **snoRNAs**: Small nucleolar RNAs **TPM**: transcripts per million **qPCR**: quantitative real-time PCR **GAPDH**: glyceraldehyde-3-phosphate dehydrogenase **GEO**: Gene Expression Omnibus **NPC**: nasopharyngeal carcinoma

## Declarations

## Acknowledgments

The skillful technical assistance of Shanghai Genergy Bio-Corporation is gratefully acknowledged.

### Competing interest

The authors declare no financial conflict of interest

### Author Contributions

All authors have read and approved of the submission of the manuscript. Conceived and designed the experiments: YQG CSJ WXT. Performed the experiments: WB YE. Analyzed the data: WB YE HY. Contributed reagents/materials/analysis tools: YQG WB YE.Wrote the paper: WB YN.

## Supporting Information

**Figure 1.**
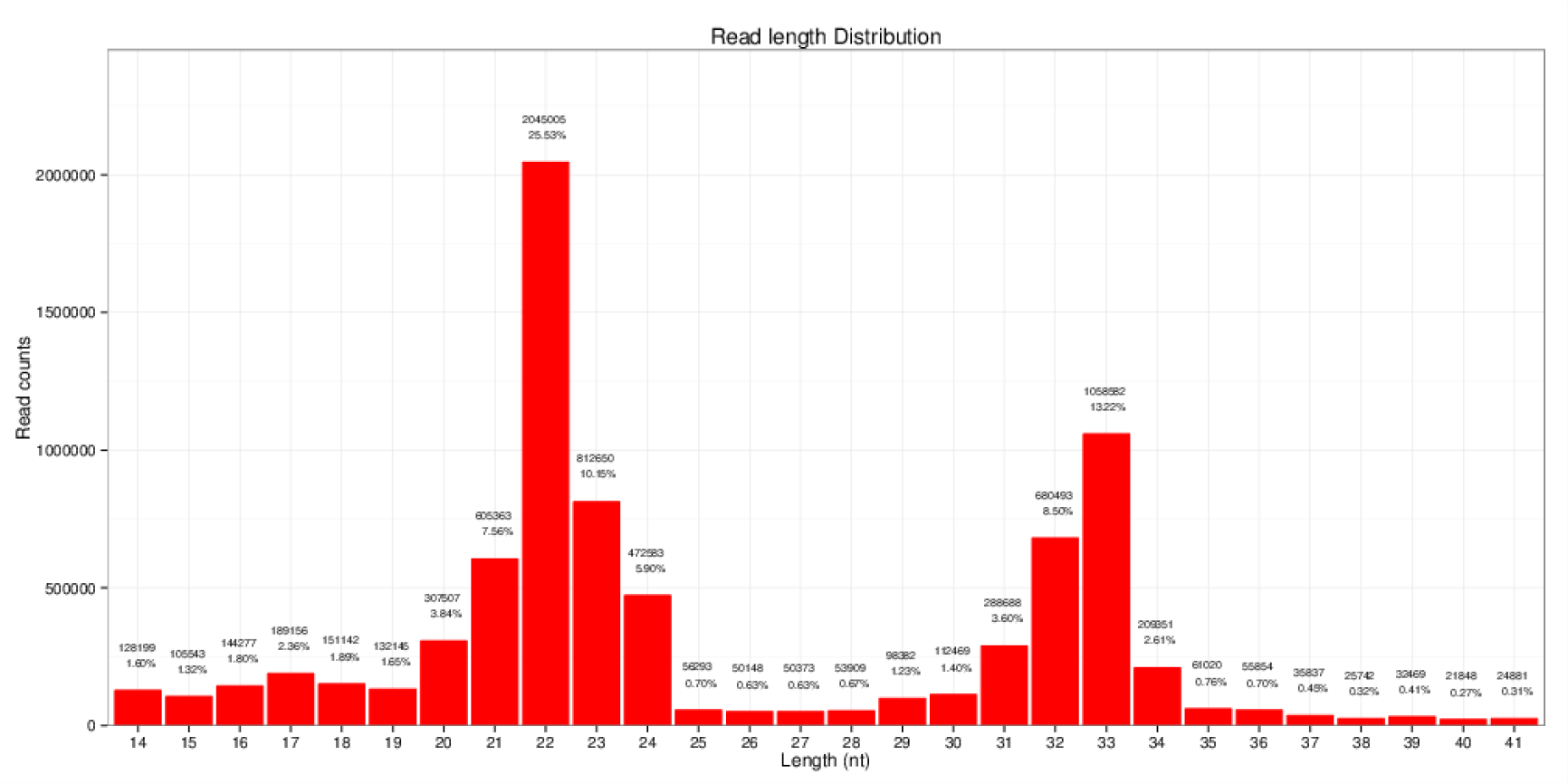
Reads length distribution statistical of S1

**Figure 2.**
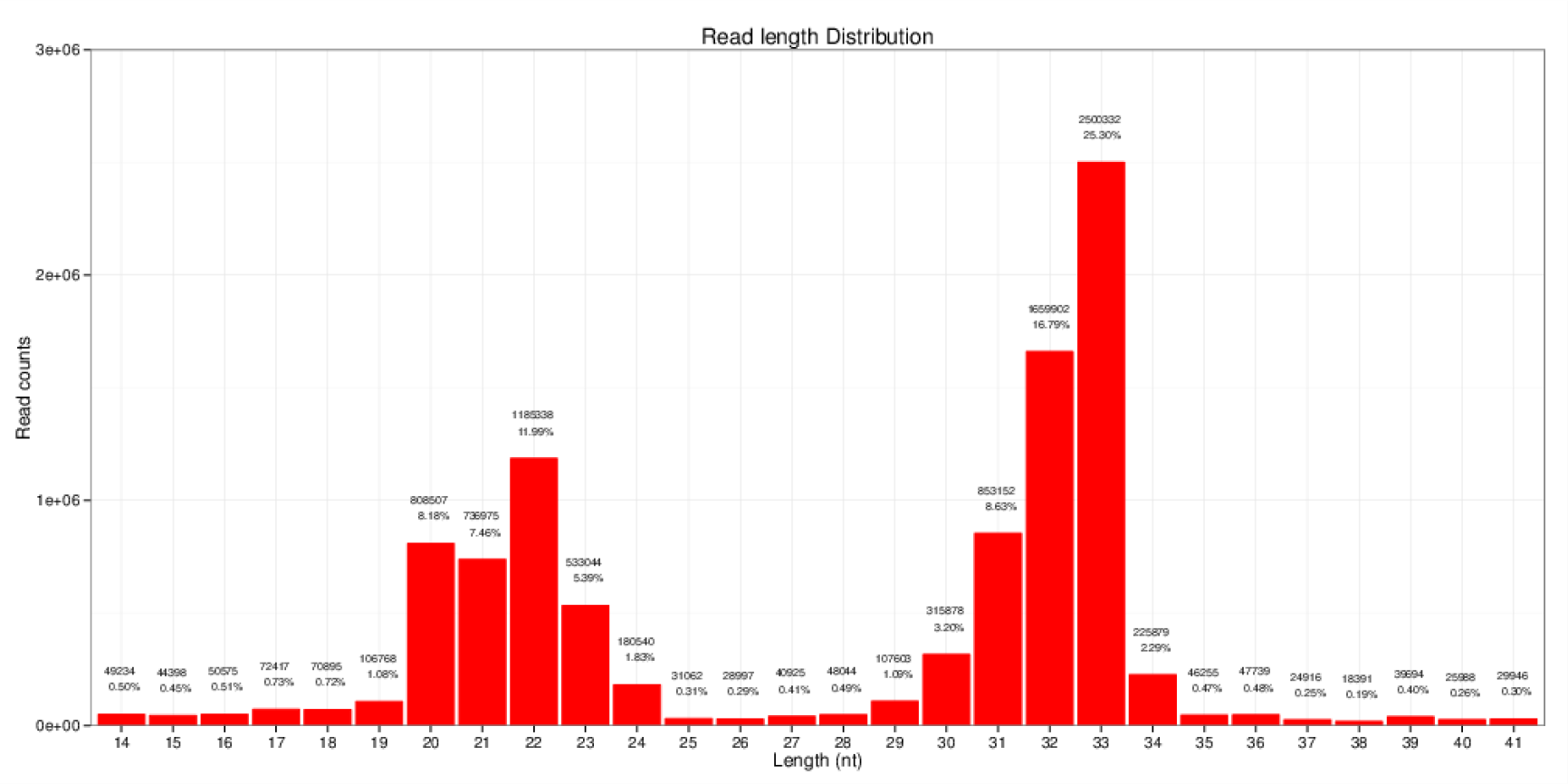
Reads length distribution statistical of S2

**Figure 3.**
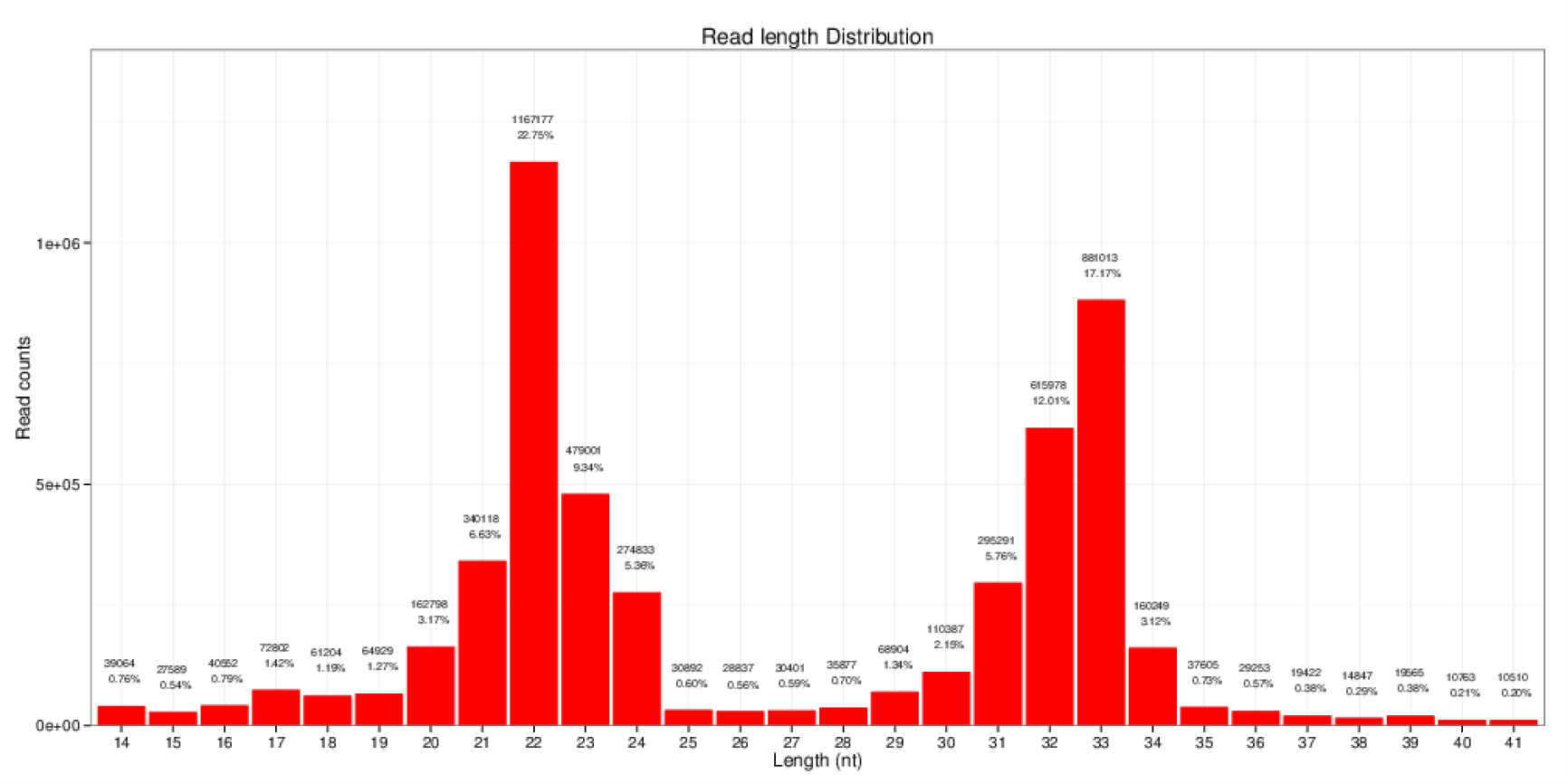
Reads length distribution statistical of S3

**Figure 4.**
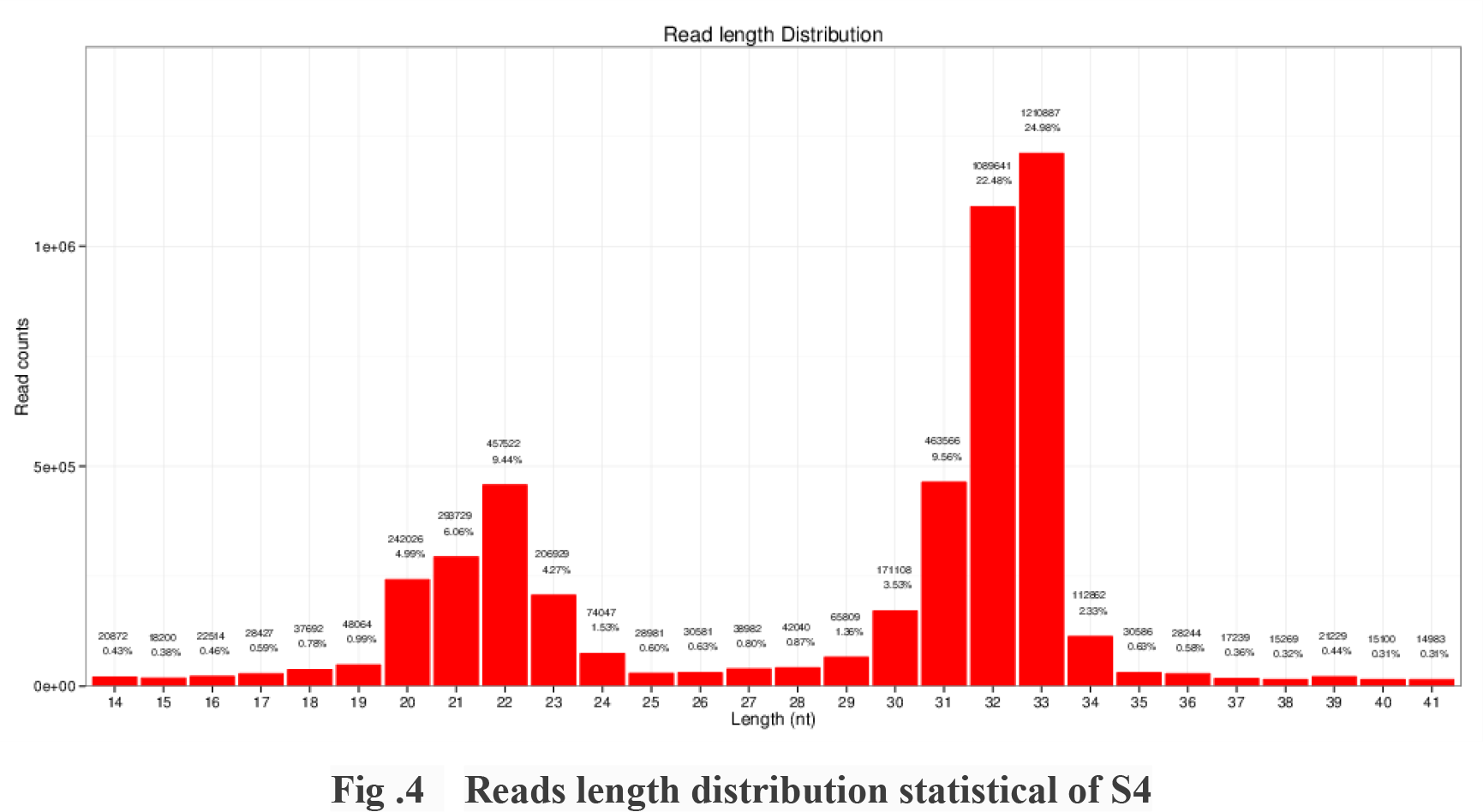
Reads length distribution statistical of S4

**Figure 5.**
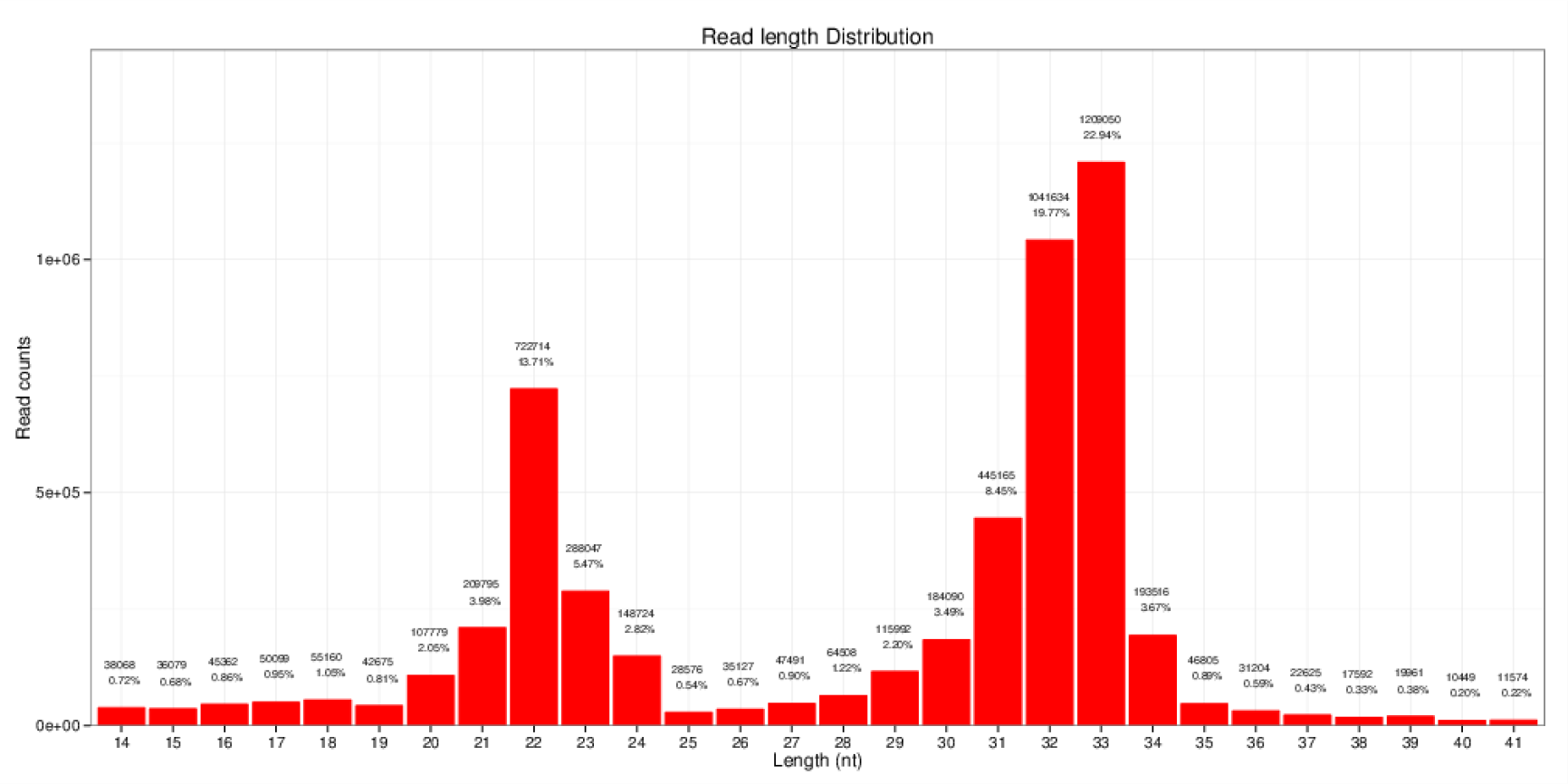
Reads length distribution statistical of S5

**Figure 6.**
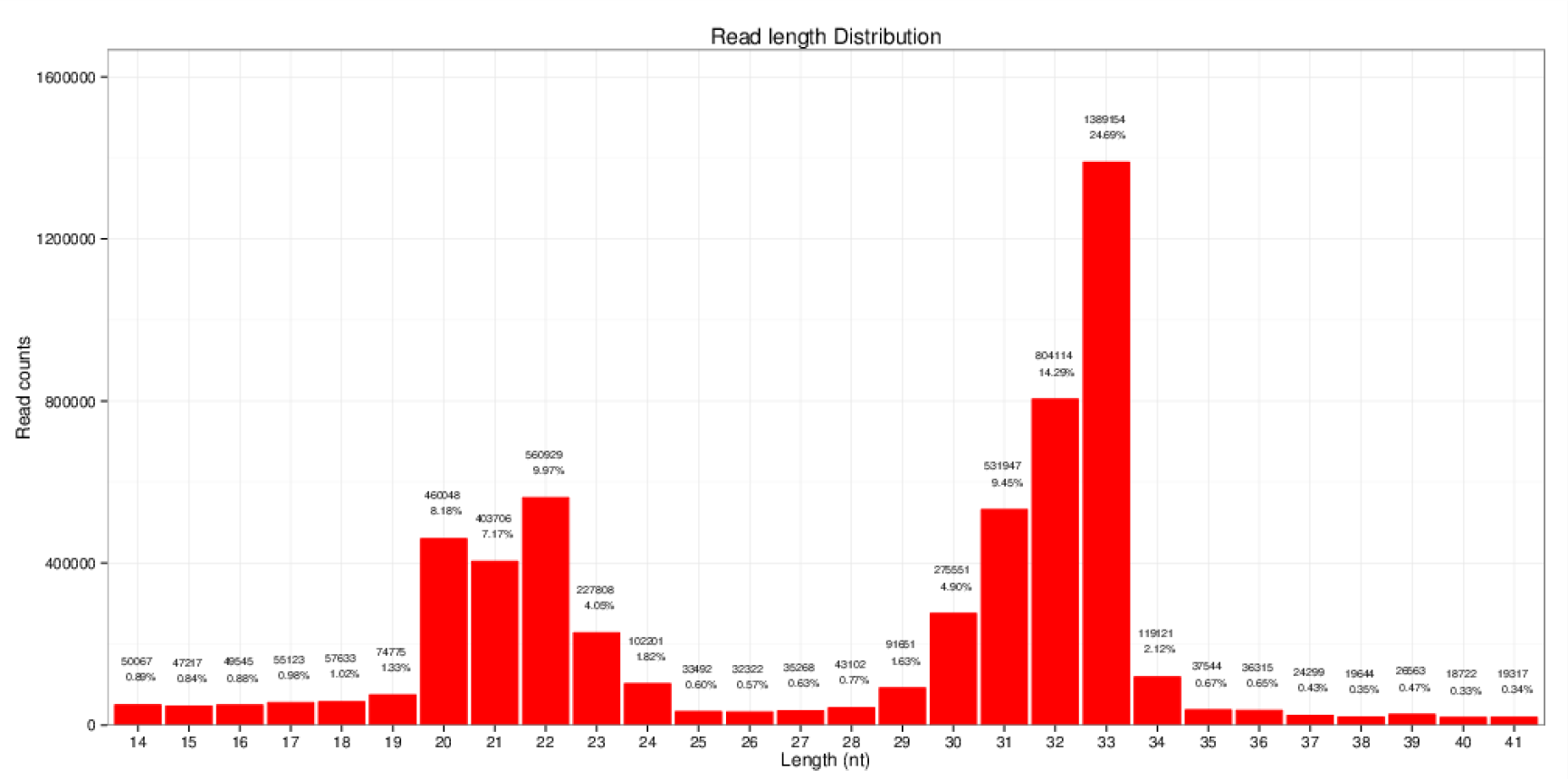
Reads length distribution statistical of S6

**Figure 7.**
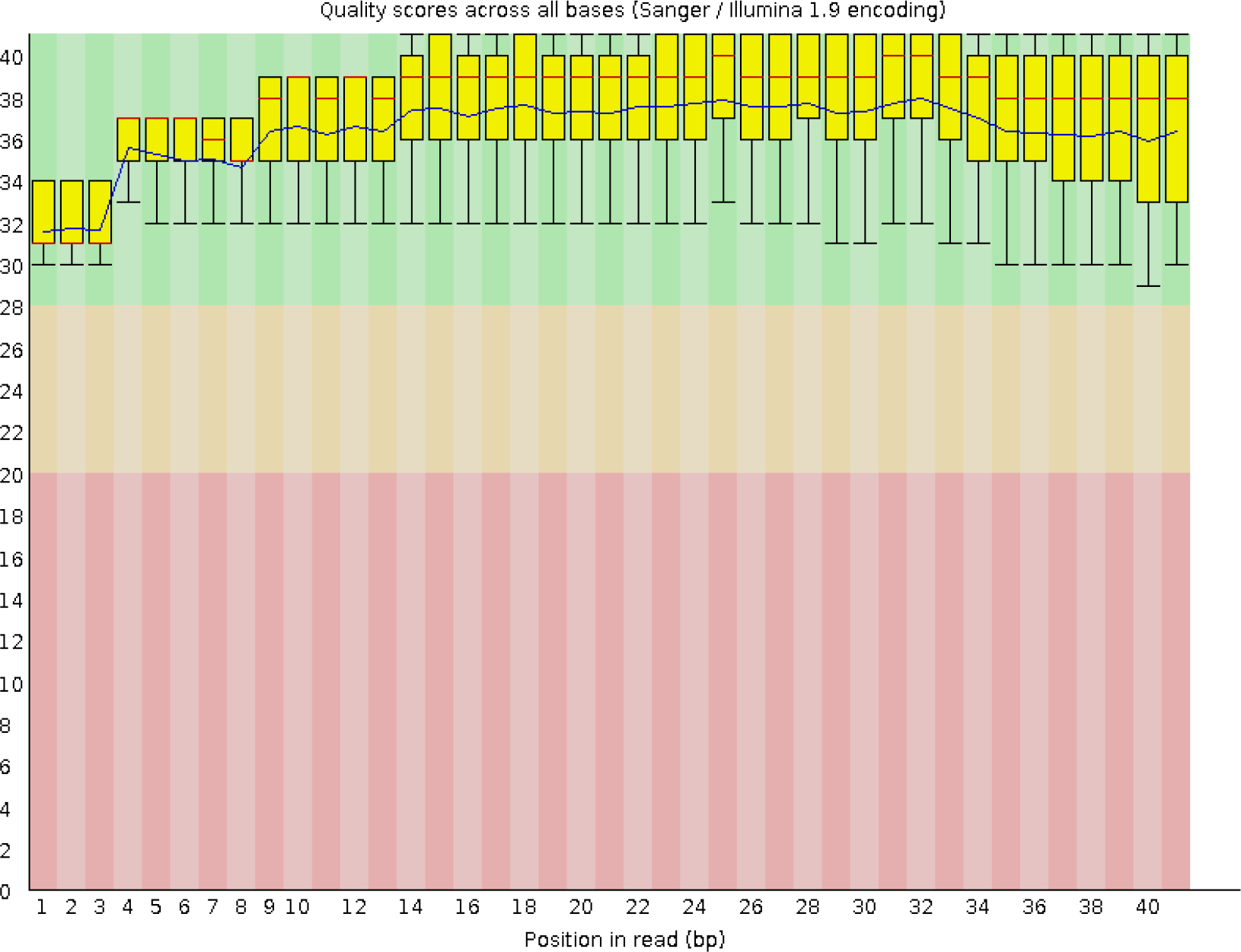
Base quality distribution in each cycle.

**Figure 8.**
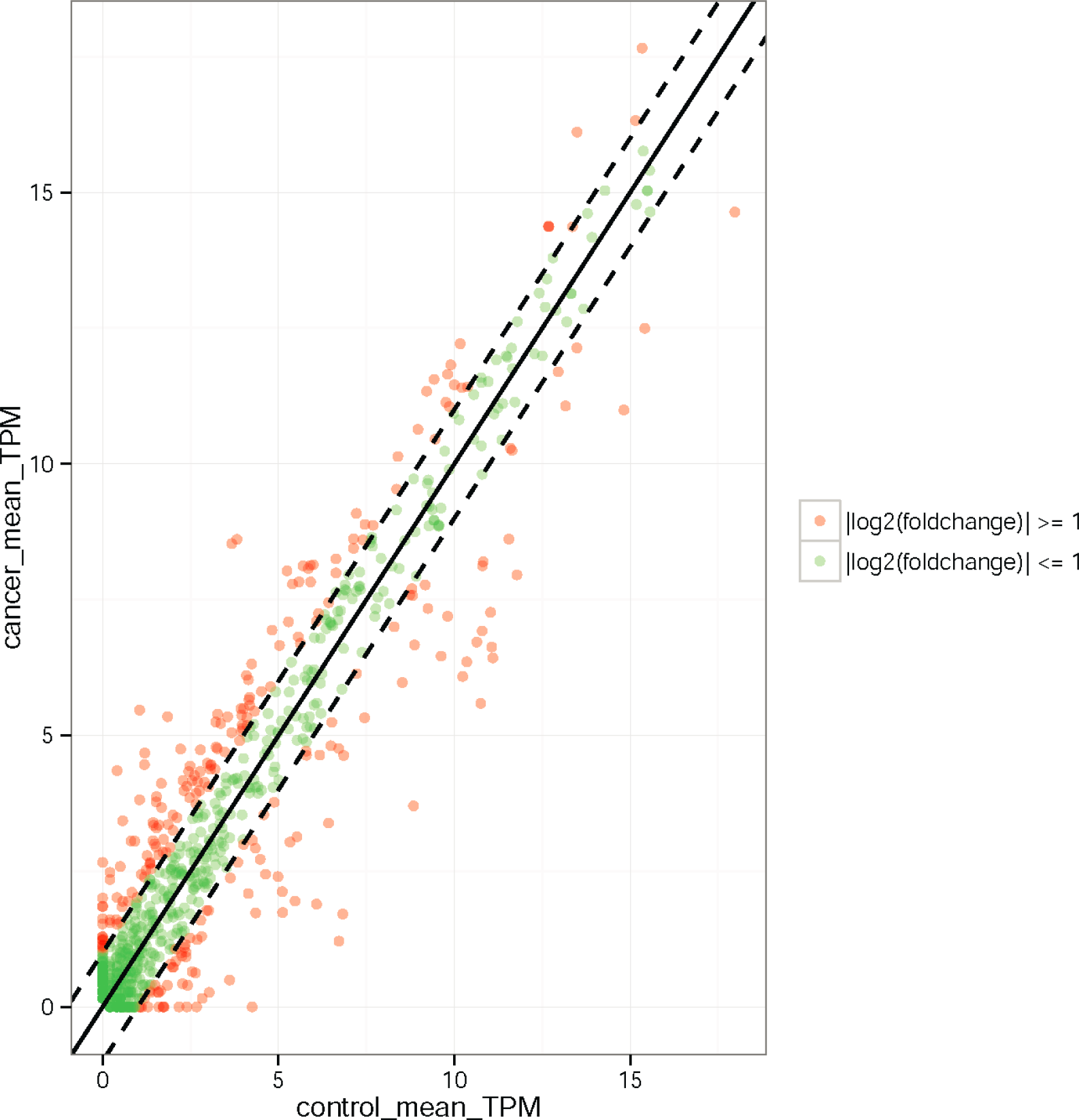
Relative expression of miRNAs in ENA and para-cancerous tissues. The *x*-and *y*-axes indicate the mean TPM expression levels of the miRNAs in each tissue. The red circles represent miRNAs with a fold change ≥ 2; green circles represent miRNAs withafold change ≤ 2 the points on the dotted line represent miRNAs witha fold change = 2. Fold changes were calculated as the mean miRNA TPM in ENA/mean miRNA TPM in para-cancerous tissues.

**Figure 9.**
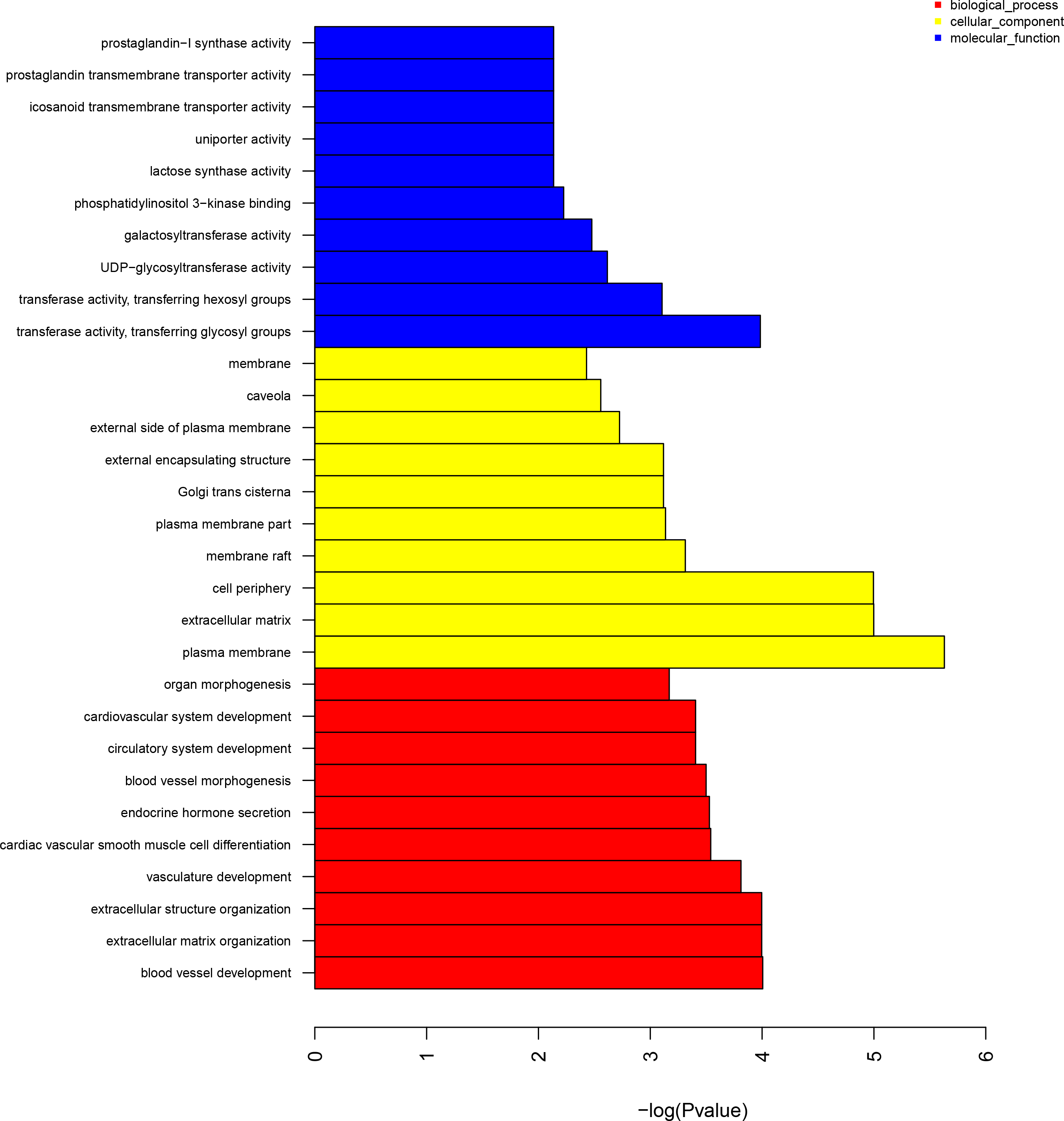
The ten most-enriched GO categories of the differentially expressed target genes of the differentially expressed miRNAs.

**Figure 10.**
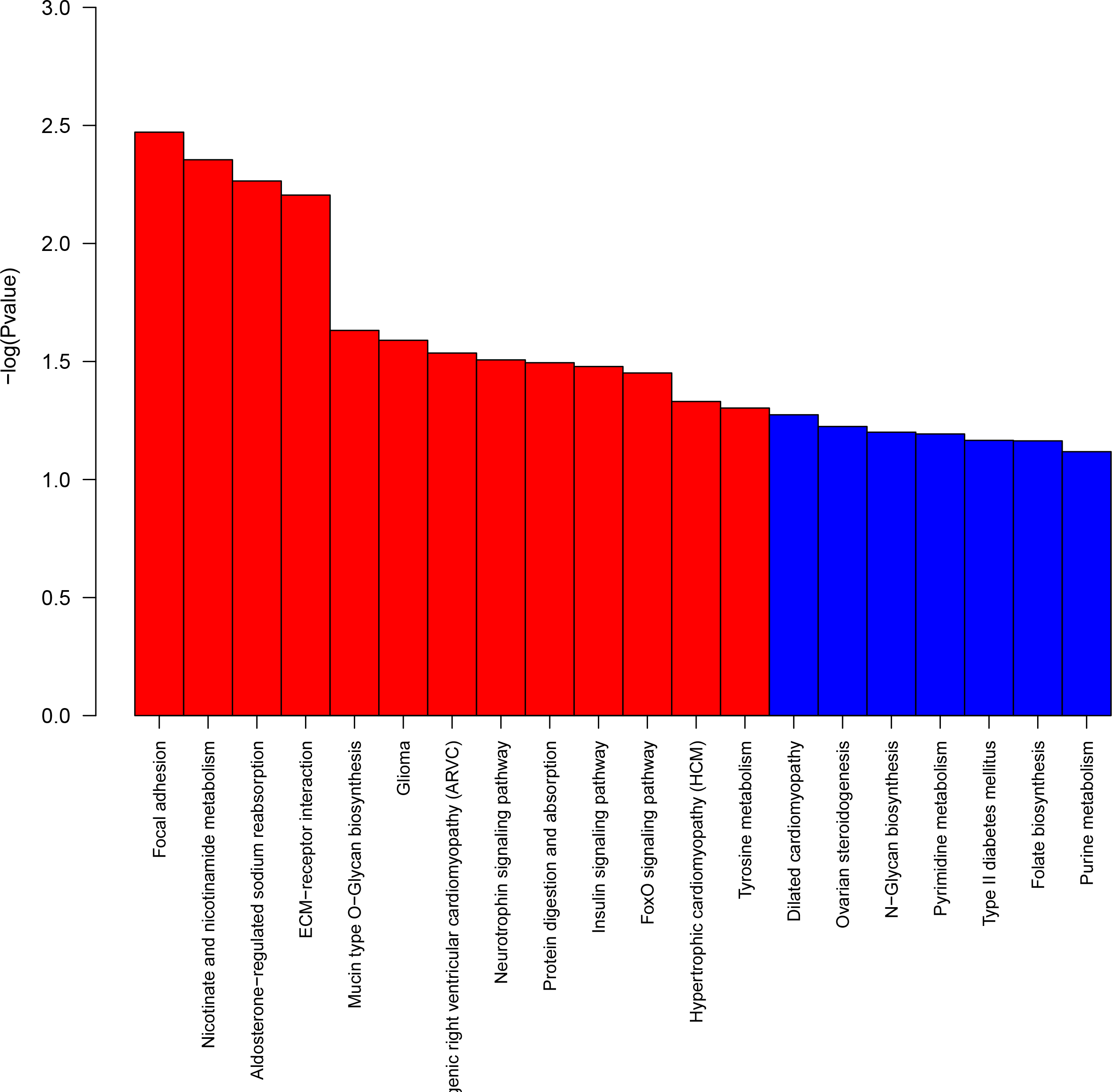
The ten most-enriched signaling pathways of the differentially expressed target genes of the differentially expressed miRNAs.

**Figure 11.**
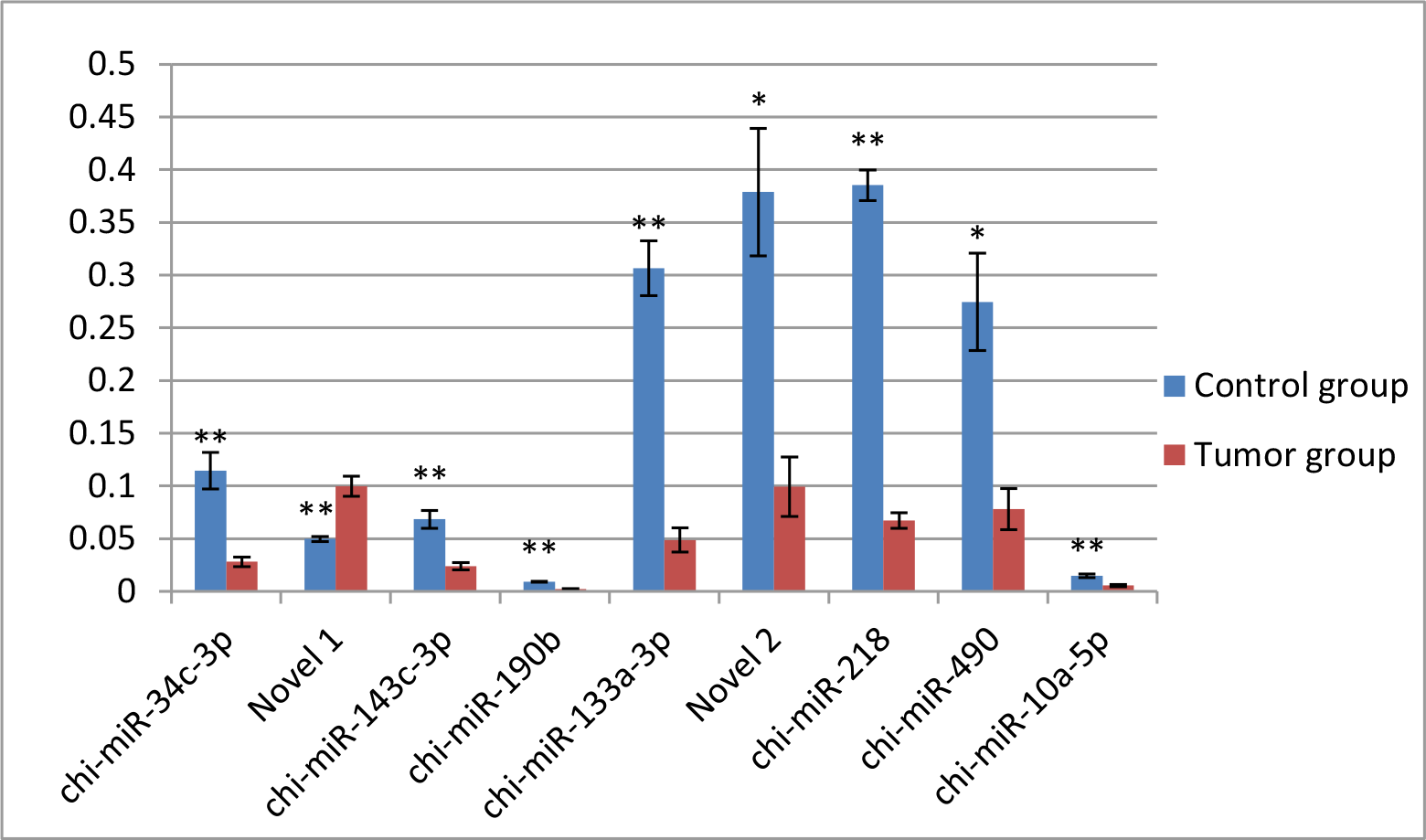
QRT-PCR validation of the identified miRNAs using Illumina sequencing technology. Real-time RT-PCR analysis of nine miRNAs in the tumour and para-carcinoma tissues from 5 goats with ENA. Relative quantification was assessed using the 2^-^ΔΔ^Cq^ method and was normalized to *U6* and *GAPDH.* 2^-^ΔΔ^Ct^ Means ±SE relative expression levels are presented. *represents *p*<0.05,** represents *p*<0.01.

Table S1. The miRNAs expressed in samples detected by Illumina sequencing. (XLSX)

Table S2. Blast results of all miRNAs against human miRNAs in miRBase v21. (XLS)

Table S3. The expression of miRNAs in ENA and para-cancerous tissues. (XLS)

Table S4. Significantly differentially expressed miRNAs in ENA.(.XLSX)

Positive FoldChange_Log2 indicates upregulation in ENA relative to the para-cancerous tissues; a negative FoldChange_Log2 indicates downregulation in ENA relative to the para-cancerous tissues; inf indicates no expression in para-cancerous tissues;-inf indicates no expression in ENA.

Table S5. Predicted target genes of the differently expressed miRNAs in ENA. (XLS)

Table S6. Differentially expressed miRNAs and their differentially expressed target genes in ENA.(XLS)

Table S7. Node attributes of the differentially expressed miRNAs and their differentially expressed target genes in ENA.(XLSX)

Table S8. Significantly enriched gene ontology categories the differentially expressed target genes of the differentially expressed miRNAs.(XLS)

Table S9. Significantly enriched signaling pathways (m=83)of gene targets of the significantly increased miRNAs in ENA. (XLS)

Table S10. Significantly enriched signaling pathways (m=89)of gene targets of the significantly reduced miRNAs in ENA.(XLS)

